# Penaeidins are a novel family of antiviral effectors against WSSV in shrimp

**DOI:** 10.1101/467571

**Authors:** Bang Xiao, Qihui Fu, Shengwen Niu, Haoyang Li, Kai Lǚ, Sheng Wang, Bin Yin, Shaoping Weng, Chaozheng Li, Jianguo He

## Abstract

Penaeidins are members of a family of key effectors with broad anti-bacterial activities in penaeid shrimp. However, the function of penaeidins in antiviral immunity is rarely reported and remains largely unknown. Herein, we uncovered that penaeidins are a novel family of antiviral effectors against white spot syndrome virus (WSSV). Firstly, RNAi *in vivo* mediated knockdown of each penaeidin from four identified penaeidins from *Litopenaeus vannamei* resulted in elevated viral loads and rendered shrimp more susceptible to WSSV, whilst the phenotype of survival rate in penaeidin-silenced shrimp can be rescued via the injection of recombinant penaeidin proteins. Moreover, pull-down assays demonstrated the conserved PEN domain of penaeidin was able to interact with WSSV structural proteins. Furthermore, we observed that colloidal gold-labeled penaeidins were located on the outer surface of the WSSV virion. By infection-blocking assay, we observed that hemocytes had lower viral infection rates in the group of WSSV preincubated with penaeidins than those of control group. Phagocytic activity analysis further showed that penaeidins were able to inhibit phagocytic activity of hemocytes against WSSV Taken together, these results suggest that penaeidins specifically binds to WSSV virion by interacting with its structural proteins, thus preventing viral infection that confers host against WSSV. In addition, dual-luciferase assay and EMSA assay demonstrated that penaeidins were regulated by Dorsal and Relish, two transcription factors of the canonical Toll and IMD pathway, respectively. To our best knowledge, this is the first report on uncovering the antiviral function of penaeidins in the innate immune system of shrimp.

**Importances:** White spot syndrome, caused by white spot syndrome virus (WSSV), is the most serious disease in shrimp aquaculture, which has long been a scourge of cultured shrimp industry. Herein, we provided some substantial evidences to indicate that penaeidins are a novel family of effectors with antiviral activity against WSSV in shrimp. Penaeidins such as BigPEN, PEN2 and PEN3 were able to interact with the outer surface of WSSV virion via binding to viral structural proteins, and thus preventing viral entry host cells. In addition, we demonstrated that the Toll and IMD signaling pathways can regulate the transcriptional expression of penaeidins, which may suggest an important role of the conserved innate signaling pathways in antiviral immunity. This is the first report of the antiviral mechanism of penaeidins in shrimp, which may provide some new insights into strategies to control WSSV infection in shrimp farms.

## Introduction

White spot syndrome virus (WSSV), which is a big dsDNA virus and considered to be the most threatening infectious pathogen in shrimp aquaculture, has caused enormous economic losses (1). WSSV infection in shrimp can cause a cumulative mortality up to 100% within 3-10 days (2). Although great progresses have been made in implications for viral prevention and control measures, including vaccination (3), immunostimulants (4), direct neutralization by antiviral proteins (5) and RNAi (6), they have not successfully restricted the uncontrolled occurrence and rapid spread of this disease in the shrimp farms. Therefore, deeper understanding the immune defense mechanism of shrimp might help to find new strategies and methods against WSSV infection.

Host innate immune system plays a significant role in protecting the organism from pathogenic invasion, particularly in invertebrates, which lacking the adaptive immunity (7, 8). Antimicrobial peptides (AMPs) are important components of the innate immune system. AMPs are originally defined as membrane-active molecules with small molecular mass (<10 kDa) that show antimicrobial activities (9). AMPs are active against a large spectrum of microorganisms: bacteria, fungi, parasites and viruses, and even against tumor cells (10). In shrimps, several families of AMPs have been identified and characterized, including Penaedins (PENs), Crustins (CRUs), anti-lipopolysaccharide factors (ALFs), Lysozymes (LYZs) and Stylicins (STYs) (11, 12). In response to viral infection, target cells can produce numerous antiviral agents to prevent the viral invasion. AMPs are one class of such agents that are typically induced in the early innate immune response (13). Direct interactions between AMPs and structural components of the virion, particularly some enveloped viruses, could be a common inhibition mechanism for destroying or destabilizing the virus and rendering it non-infectious (14). In mammals, it has been reported that cathelicidins prevent the Influenza virus to infect the host cell by direct binding to the virus and destroying its membrane (15). Indolicidin, an AMP, inactivates human immuno-deficiency virus type 1 (HIV-1) by damaging the virion membrane (16). In addition, α-defensin peptides can neutralize herpes simplex virus 1 (HSV-1) by binding to natural viral glycoproteins (17). In shrimps, so far, the ALFs and LYZs have been demonstrated to exhibit anti-WSSV activities by binding to WSSV structural proteins (18). Until now, although penaeidins have not be reported to inhibit any viruses, the mRNA levels of PmPEN5 from *Penaeus monodon* were significantly induced when an infection with WSSV, suggesting its possible role in shrimp antiviral immunity (19). However, whether penaeidins are an important class of antiviral effectors against WSSV, and if so, how the antiviral mechanisms of penaeidins execute deserve to be studied and explored.

Penaeidins belong to an AMP family initially characterized from the shrimp *Litopenaeus vannamei* and play a significant role in antibacterial immunity (20). Penaeidins are unique cationic molecules that consist of an N-terminal proline-rich region (PRR) and a C-terminal cysteine-rich region (CRR) within six conserved cysteine residues forming three disulfide bonds (21). Penaeidins can be classified into four distinct subgroups: PEN2, PEN3, PEN4 and PEN5 (as PEN1 turned out to be the variant of PEN2) based on amino acid sequence comparisons and the position of specific amino acids (22). It has been reported that usually more than one subgroup was found in a penaeid shrimp. For example, three penaeidins subgroups (PEN2, PEN3 and PEN4), were identified in *L. vannamei* and *Litopenaeus setiferus* (23), whilst two subgroups of penaeidins, PEN3 and PEN5, were found in *Fenneropenaeus chinensis* (24) and *P. monodon* (25). Interestingly, penaeidins are members of a family of key effectors with broad anti-bacterial activities only found in penaeid shrimps. However, the function of penaeidins in antiviral immunity is rarely reported and remains largely unknown.

In this study, we obtained a new PEN cDNA from the *L. vannamei*, and designated it as BigPEN according to its additional repeat (RPT) region and high molecular weight. All four subgroups of penaeidins from *L. vannamei* including BigPEN and previously identified PEN2, PEN3 and PEN4 were chosen to explore their function during WSSV infection. Our results showed that they were able to bind to the surface of WSSV virion and its structural proteins. In addition, each recombinant penaeidin inhibited phagocytic activity of hemocytes against WSSV, and attenuated the ability of WSSV to infect hemocytes. Moreover, the Toll and IMD pathways (canonical NF-κB pathways) were demonstrated to regulate the production of four subgroups of penaeidins. Taken together, we provided the first evidence that the NF-κB pathway controlled penaeidins is a novel class of antiviral effectors against WSSV in shrimp.

## Results

### Penaeidins were strongly upregulated *in vivo* after WSSV infection

Penaeidins have been previously identified as AMPs with significant antibacterial and antifungal activities, but no antiviral activity has been reported. To explore whether penaeidins have any antiviral roles in defense against WSSV, a major viral pathogen in shrimp, we firstly searched the EST sequences homologous to known penaeidin proteins from our transcriptome in *L. vannamei* (26), and obtained a new paralog. We cloned the full-length cDNA sequences of the new paralog by using rapid amplification cDNA ends (RACE)-PCR method, and we subsequently designated it as BigPEN as it containing an additional repeat (RPT) region and its high molecular weight (29.22 kDa). Until now, a total of four penaeidins including the newly cloned BigPEN and previously identified PEN2, PEN3 and PEN4 were presented in shrimp *L. vannamei*, which were clustered into three major groups (Fig. 1C). As shown in Fig. 1D, BigPEN has an additional RPT domain compared with PEN2, PEN3 and PEN4, all of which contain a conserved PEN domain consisted of a proline-rich region (PRR) and a C-terminal cysteine-rich region (CRR). Of note, penaeidins are only found in penaeid shrimps. To understand the function of penaeidin family during WSSV infection more comprehensively and thoroughly, BigPEN, together with previously identified PEN2, PEN3 and PEN4, were all chosen to address the tissue distributions in healthy shrimps and time-course expression patterns in viral challenged shrimps. By quantitative reverse transcription PCR (qRT-PCR), we observed that the four penaeidins were mainly expressed in hemocytes of naïve (uninfected) shrimp (Fig. 1A), and thus the hemocyte was used to be the target tissue for following analysis. In regard of penaeidins as conventional AMPs, *Vibrio parahaemolyticus* was chosen to be a control pathogen to compare the expression levels of penaeidins after WSSV and this bacterial infection. We found that both of pathogens could markedly induce the expression of all four penaeidins in the early stage of infections in hemocytes (Fig. 1B). In particular, the degrees of upregulation of BigPEN and PEN2 during 4-12 h after *V. parahaemolyticus* were more than those of the treatment of WSSV, whereas they displayed different expression profiles during 24-48 h with a slight upregulation or downregulation. The PEN3 showed increased expression patterns during 4-8 h after WSSV infection, but suppressed expression patterns during 12-48 h. The transcriptional levels of PEN4 in response to WSSV challenge were sharply up-regulated during 4-24 h, but down-regulated at 36 h (Fig. 1B). Taken together, these results suggested that the induced penaeidins might participate in the immune response against pathogenic encroachment in *L. vannamei*.

**Figure 1.**
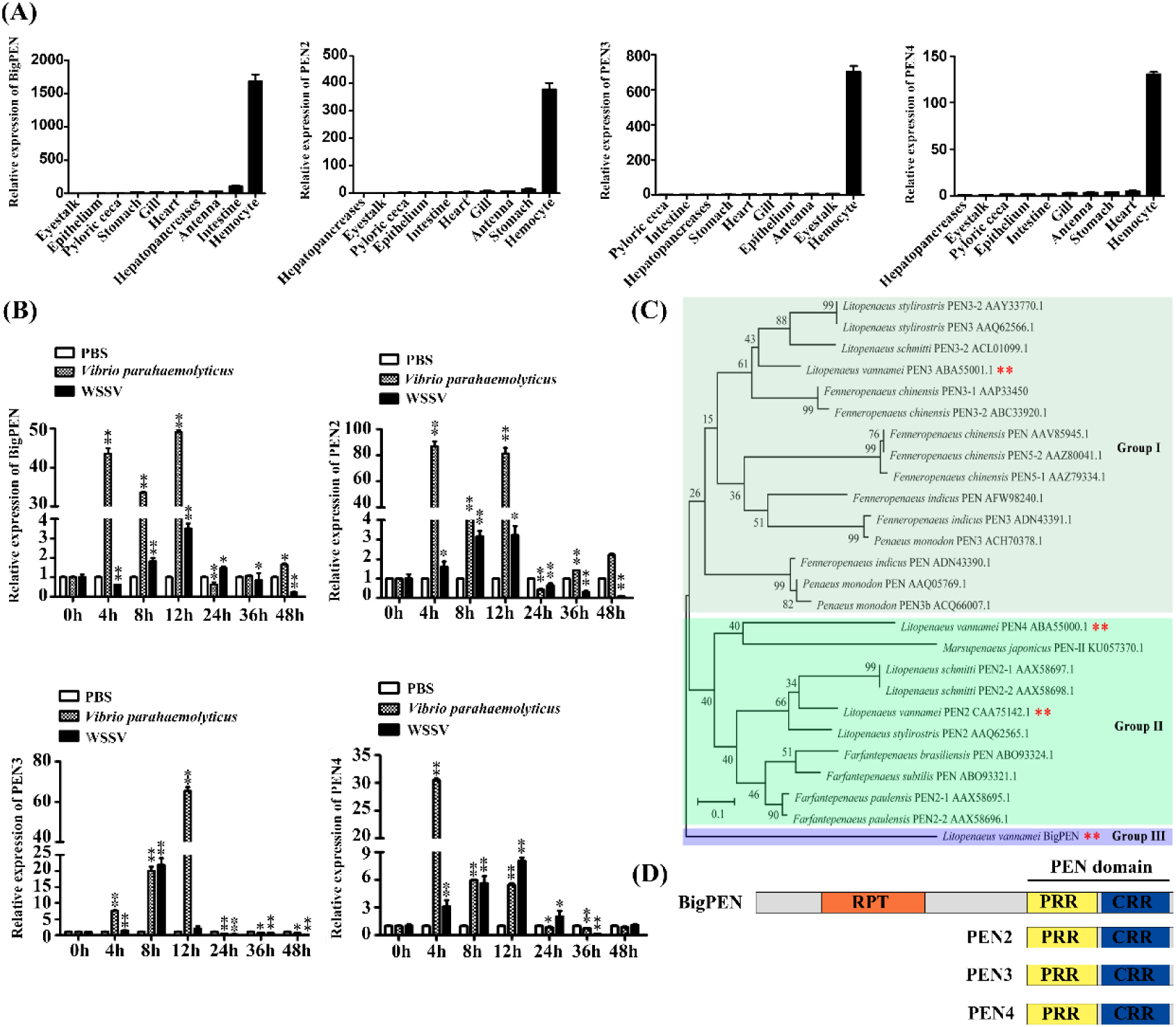
Penaeidins were strongly induced in response to WSSV infection. (A) Transcriptional levels of BigPEN, PEN2, PEN3 and PEN4 in different tissues of healthy shrimps were analyzed by quantitative RT-PCR. *L. vannamei* EF-1α was used as an internal control and the data were shown as means ± SD of triμlicate assays. (B) Expression profiles of BigPEN, PEN2, PEN3 and PEN4 in hemocytes from WSSV or *V. parahaemolyticus* or PBS (as a control) challenged shrimps. Quantitative RT-PCR was performed in triμlicate for each sample. Expression values were normalized to those of EF-1α using the Livak (2^-–ΔΔCT^) method and the data were provided as means ± SD of triplicate assays. The statistical significance was calculated using Student’s t-test (^**^ *p* < 0.01 and ^*^ *p* < 0.05). (C) Phylogenetic tree was constructed using amino acid sequences of PEN domains from different penaeidins. The GenBank accession numbers are showed after scientific names of their species. (D) Architecture diagrams of BigPEN, PEN2, PEN3 and PEN4. Penaeidin domain and different regions are shown with distinct colors.

### Penaeidins restricted WSSV replication *in vivo*

To obtain the function of penaeidins during WSSV infection, RNAi *in vivo* combined with the injection of recombinant penaeidin proteins were performed. We designed and synthesized different dsRNAs namely dsRNA-BigPEN, dsRNA-PEN2, dsRNA-PEN3 and dsRNA-PEN4, which can specifically target mRNAs of BigPEN, PEN2, PEN3 and PEN4, respectively. As shown in Fig. 2A, the mRNA levels of each penaeidin could be effectively suppressed by corresponding gene-specific dsRNA. After the knockdown of penaeidins, the shrimps were infected with WSSV by intramuscular injection, and the viral loads (WSSV copies) in the each penaeidin-silenced shrimp was determined by absolute quantitative PCR (absolute q-PCR) at 48 hours post infection (hpi). We observed that a greater number of each penaeidin silenced shrimps exhibited higher quantities of viral titers in muscles when compared to control shrimps (Fig. 2B). To further demonstrate the anti-WSSV role of penaeidins, experiments of RNAi *in vivo* plus recombinant penaeidin proteins were performed. We observed that shrimps with the co-injection of recombinant penaeidin plus dsRNA had reduced viral replication levels compared to the control group (Fig. 2C). These results strongly indicate that all four penaeidins can inhibit WSSV replication *in vivo*. To investigate whether the changes of each penaeidin mediated viral replication levels *in vivo* are implicated with resistance or tolerance to WSSV, survival rate experiment was performed and recorded. We observed that only knockdown of PEN2 resulted in significantly lower survival rate than the GFP dsRNA control group (*P* = 0.0135 < 0.05) (Fig. 2E). Nevertheless, other shrimps with knockdown of BigPEN, PEN3 or PEN4 still showed reduced survival rates to some extent, despite of no significant difference in the statistical analysis compared with corresponding control group (Fig. 2D, 2F and 2G). It is noteworthy that each recombinant penaeidin protein can notably confer shrimps more resistance to WSSV (*P* < 0.01) (Fig. 2D-2G). This above phenomenon could be explained by that the effect of knockdown of single penaeidin via RNA *in vivo* might be replenished by other ones or effectors in an unidentified mechanism, whereas injection of recombinant penaeidin not only rescues the silenced one but also confer shrimps more protection from WSSV. In summary, these results convincingly demonstrate that penaeidins are a class of critical antiviral factors against WSSV *in vivo*.

**Figure 2.**
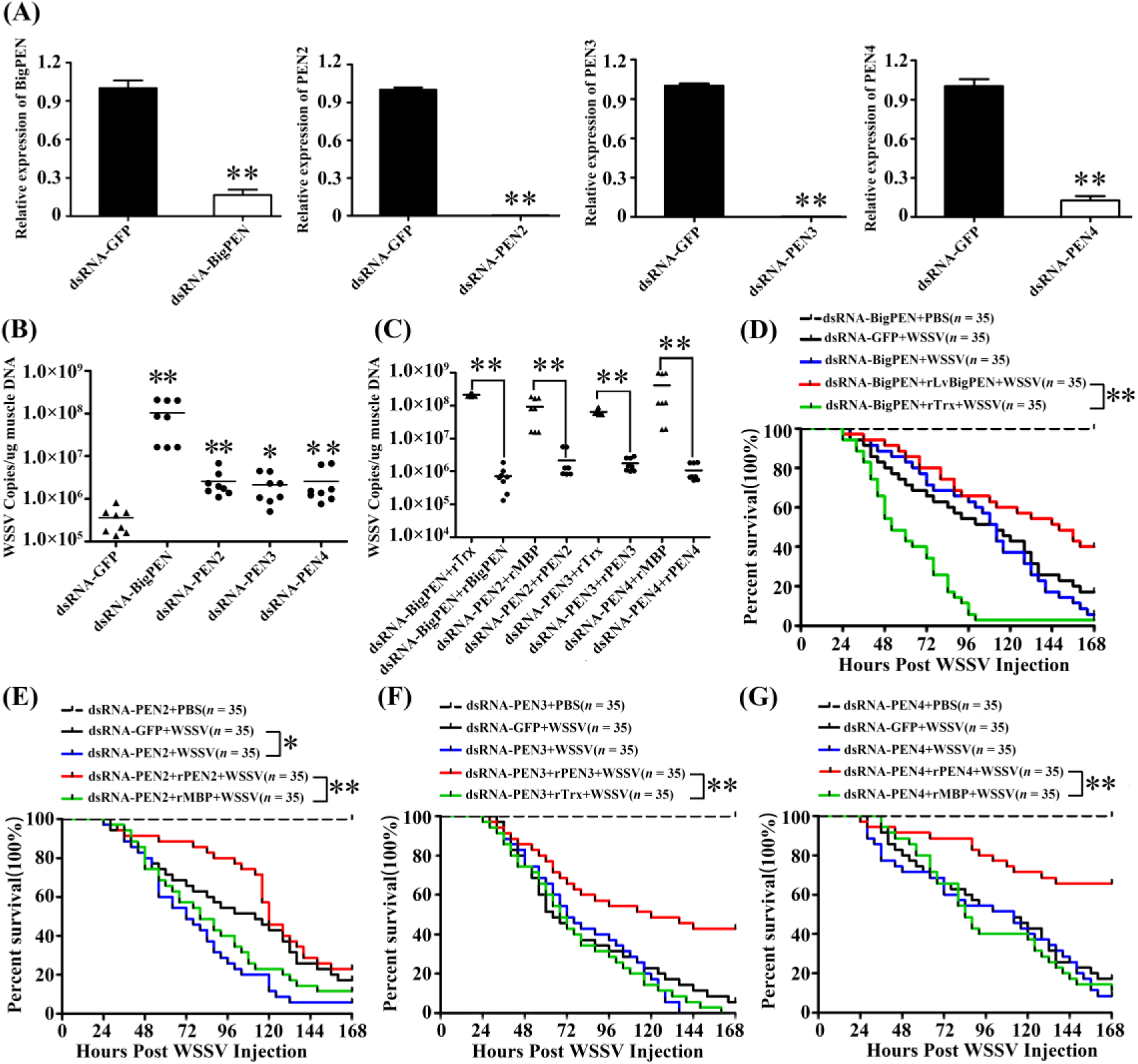
Penaeidins possessed potent antiviral activities against WSSV. (A) Quantitative RT-PCR analysis of the silencing efficiencies of BigPEN, PEN2, PEN3 and PEN4, the internal control was EF-1α. Samples were taken at 48 h after injection with gene specific dsRNA or GFP dsRNA. (B) The quantity of WSSV copies in muscles from each individual shrimp in five different groups were detected by absolute quantitative PCR. After 48 h of WSSV infection, eight shrimps were chosen to be detected WSSV copies in each group. Differences between experiment groups and control group (GFP dsRNA) were analyzed using Student’s t-test (^**^ *p* < 0.01 and ^*^ *p* < 0.05). (C) WSSV copies in muscles of each individual shrimp from four groups were detected by absolute quantitative PCR. After 48 h of dsRNA injection, the shrimps were injected with WSSV premixed with the purified rBigPEN, rPEN2, rPEN3 or rPEN4, respectively. Injections of the same amount of a mixture of WSSV with Trx or MBP proteins were used as controls. The muscles from each group (8 shrimps) were sampled for absolute quantitative PCR to detect WSSV loads at 48 h post infection. Differences between experiment groups and control group (GFP dsRNA) were analyzed using Student’s t-test (^**^ *p* < 0.01 and ^*^ *p* < 0.05). (D-G) The survival rates of WSSV infected shrimps with knockdown of penaeidins including BigPEN (D), PEN2 (E), PEN3 (F) or PEN4 (G). Meanwhile, a series of experiments by co-injection of purified recombinant penaeidins to rescue the knockdown of each penaeidin during WSSV infection were performed. The death of shrimp was recorded at every 4 h for survival rates analysis by the Kaμlan-Meier method (^**^ *P* < 0.01). All experiments were performed three times with similar results.

### Penaeidins bound to WSSV structural proteins via their conserved PEN domains

Direct interaction of antiviral factors with viral proteins has been long postulated as a generally antiviral mechanism, especially for enveloped viruses (27, 28). To clarify the possible mechanism of penaeidins against WSSV, pull-down assay was performed to detect whether penaeidin proteins could interact with WSSV structural proteins. Because BigPEN contained an additional N-terminal RPT domain and conserved C-terminal PEN domain consisted of a PRR and a CRR (Fig. 1D, Fig. 3A), the full-length (BigPEN-FL), the RPT domain (BigPEN-R) and the C-terminal PEN domain (BigPEN-PEN) with His tag were expressed and purified (Fig. 3B). Several envelope and tegument proteins of WSSV including VP19, VP24, VP26, VP28 and VP16 with GST tag were chosen to expressed and purified (Fig. 3C). In the GST pull-down assays, we observed that GST tagged viral proteins including VP26, VP28 and VP16 precipitated BigPEN-FL by SDS–PAGE with coomassie blue staining (Fig. 3D upper panel, lanes 4, 5 and 6), and further confirmed this result by western blotting with His tag antibody (Fig. 3D down panel). In the His tagged BigPEN-FL pull-down assays with five WSSV structural proteins (GST tag), we obtained an identical result that BigPEN-FL precipitated VP26, VP28 and VP16 (Fig. 3E). To further verify which domain of BigPEN-FL was able to interact with WSSV structural proteins, two separate domains including BigPEN-R and BigPEN-PEN were used to pull-down assays, respectively. In both of GST pull-down assays and His pull-down assays, we observed that BigPEN-PEN, but not BigPEN-R, was able to interact with VP26, VP28 and VP16 (Fig. 3F-3I). Collectively, these results strongly suggest that C-terminal PEN domain of BigPEN can interact with WSSV structural proteins VP26, VP28 and VP16 (Fig. 3J).

**Figure 3.**
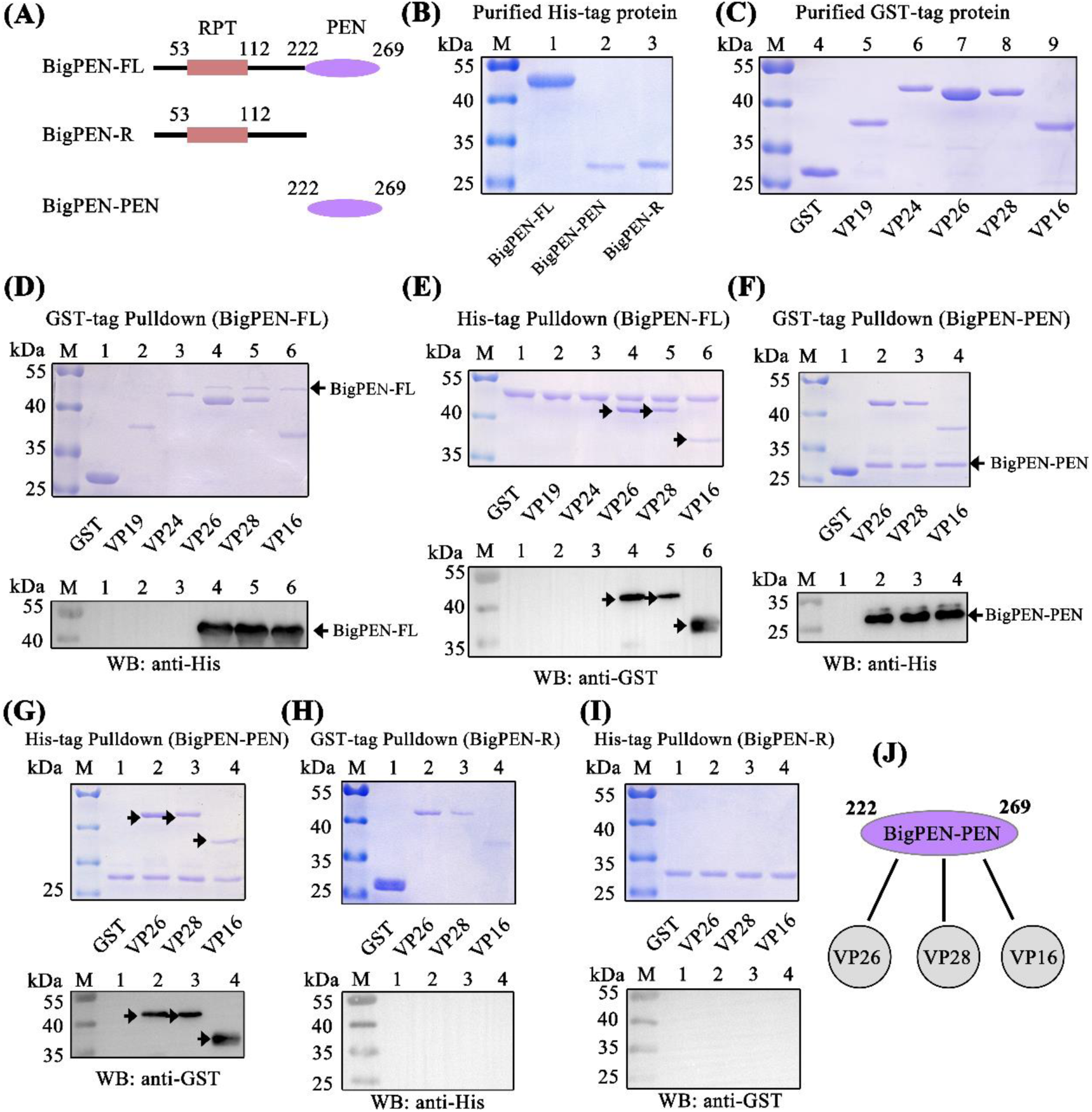
The PEN domain of BigPEN interacted with WSSV structural proteins. (A) Domain architecture of BigPEN. Three plasmids with His tag including the full-length (BigPEN-FL), the RPT domain (BigPEN-R) and the PEN domain (BigPEN-PEN) of BigPEN were generated. The red and blue represent the RPT domain and PEN domain of BigPEN, respectively. (B) Recombination expression and purification of His-tagged BigPEN-FL, BigPEN-R and BigPEN-PEN. The purified proteins were analyzed using SDS-PAGE and stained with Coomassie blue. (C) Recombination expression and purification of GST, GST-tagged VP19, VP24, VP26, VP28 and VP16. (D-E) GST-pulldown (D) and His-pulldown (E) assays to detect the interaction between BigPEN-FL with VP19, VP24, VP26, VP28 and VP16, BigPEN-FL could bind to VP26, VP28 and VP16, the analysis using 12.5% SDS-PAGE by staining with Coomassie blue and Western-blot, GST tag was performed as a control. (F-G) GST-pulldown (F) and His-pulldown (G) assays showed that BigPEN-PEN could bind to VP26, VP28 and VP16. The results were detected using 12.5% SDS-PAGE by staining with Coomassie blue and Western-blot. (H-I) GST-pulldown (H) and His-pulldown (I) assays demonstrated that RPT domain of BigPEN (BigPEN-R) can’t interact with the five WSSV structural proteins. (J) Schematic illustration of the BigPEN-PEN interacting with VP26, VP28 and VP16. All the experiments were repeated three times.

Since the C-terminal PEN domain of BigPEN showed high conservation to PEN domains of PEN2, PEN3 and PEN4, which inspired us whether PEN domains of PEN2, PEN3 and PEN4 were also able to interact with some viral structural proteins. To address this, GST pull-down assays and His pull-down assays were performed to explore the possible interaction between PEN2, PEN3 and PEN4 and above five viral proteins. In contrast to BigPEN-FL, the three penaeidins of PEN2, PEN3 and PEN4 only contained the conserved PEN domains (Fig. 4A). For an unfavorable reason, the His tagged PEN2 and PEN4 proteins failed to be achieved, and thus MBP tagged proteins was instead expressed and purified (Fig. 4B). In the GST pull-down assays, we found that only VP24 was enriched with PEN2 (Fig. 4C upper panel, lane 3), and an identical result was observed by western blotting (Fig. 4C down panel, lane 3). Likewise, in the MBP tagged PEN2 pull-down assays with GST tagged viral proteins, we observed that PEN2 was able to interact with VP24, but not other tested viral proteins (Fig. 4D). By a similar method, PEN3 was demonstrated to specially bind to VP26, but not other tested viral proteins (Fig. 4E-4F). Unexpectedly, PEN4 did not interact with the five tested viral proteins VP19, VP24, VP26, VP28 or VP16 (Fig. 4G). Taken together, these results demonstrated that PEN2 was able to interact with VP24, and PEN3 was able to interact with VP26 (Fig. 4H).

**Figure 4.**
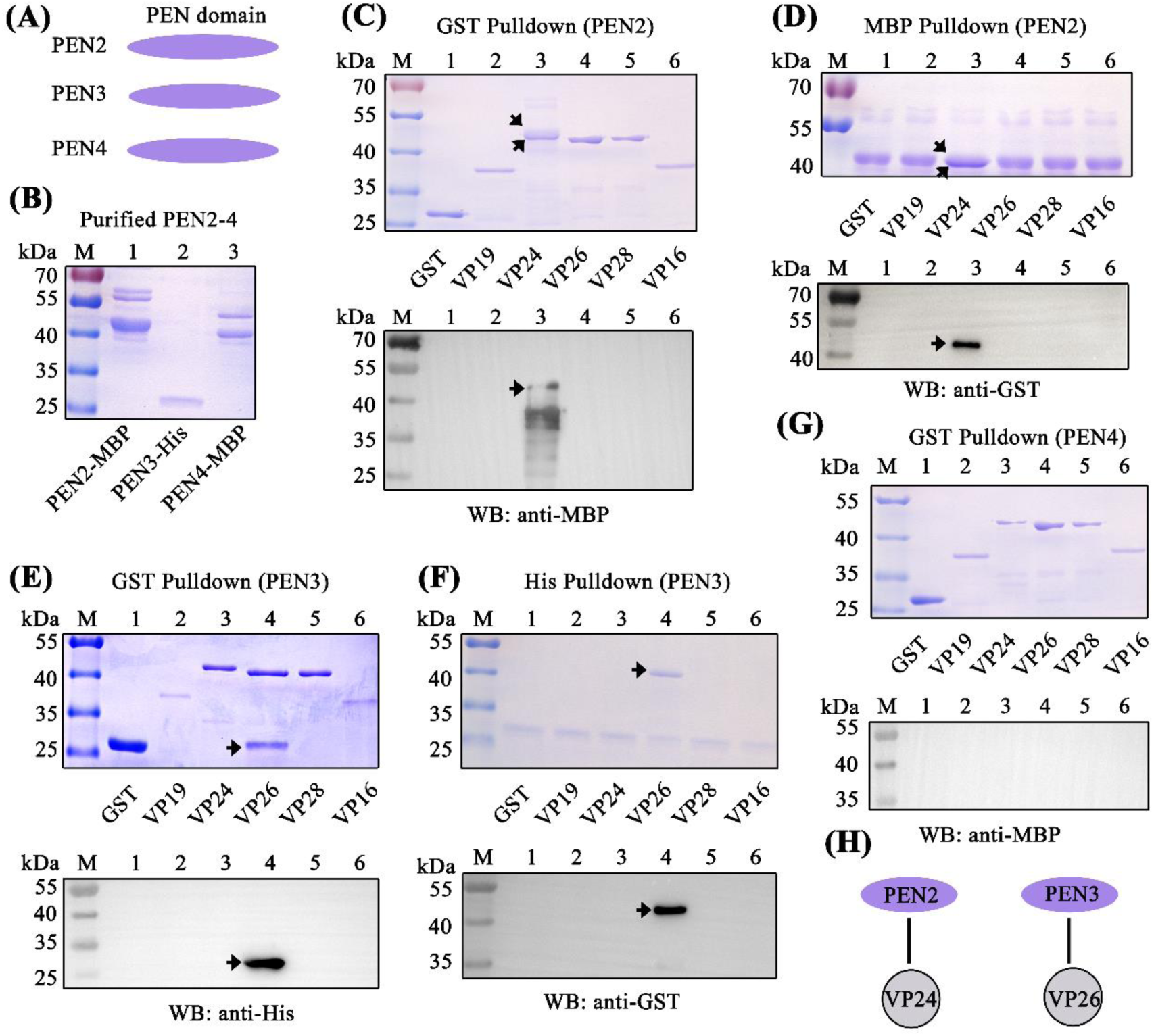
PEN2, PEN3 and PEN4 interacted with WSSV structural proteins. (A) Domain architecture of PEN2, PEN3 and PEN4. (B) Recombination expression and purification of MBP-tagged PEN2 and PEN4, and His-tagged PEN3. (C-D) GST-pulldown (C) and MBP-pulldown (D) assays to detect the interaction between PEN2 with VP19, VP24, VP26, VP28 and VP16. PEN2 could bind to VP24, which are showed using 12.5% SDS-PAGE by staining with Coomassie blue and Western-blot. (E-F) GST-pulldown (E) and His-pulldown (F) assays showed that PEN3 could bind to VP26. The assays were detected using 12.5% SDS-PAGE by staining with Coomassie blue and Western-blot, GST tag was performed as a control. (G) GST-pulldown assay showed that PEN4 can’t interact with the five WSSV structural proteins. (H) Schematic illustrations of the PEN2 interacting with VP24, and the PEN3 interacting with VP26, respectively. All the experiments were repeated three times.

### Penaeidins inhibited WSSV to infect hemocytes

The successful infection is required for virus entry into host cells. To explore whether the interaction between penaeidins with WSSV protein could inhibit the entry of WSSV into hemocyte, infection-blocking assays were performed. The results showed that each recombinant protein including rBigPEN-FL, rBigPEN-PEN, rPEN2, rPEN3 or rPEN4 was able to inhibit WSSV attachment and penetration to hemocytes *in vitro* (Fig. 5A). The infection rates of hemocytes by WSSV were then calculated, and the purified Trx tag was used as a control and set to 100%. Compared to control, the infection rates of hemocytes were remarkably suppressed in the experimental groups infected with WSSV by preincubation with rBigPEN-FL (28.80%), rBigPEN-PEN (41.31%), rPEN2 (35.89%), rPEN3 (34.38%) and rPEN4 (36.87%) (Fig. 5B). These results strongly suggested that the preincubation of penaeidins with WSSV can effectively inhibit virus entry into host hemocytes. To investigate whether penaeidins were able to interact with WSSV virion, an experiment by colloidal gold electron microscopy was performed. We observed that each colloidal gold-labeled penaeidin was located on the outer surface of WSSV (Fig. 5C), which was consistent with the above results that penaeidins were able to interact with one or more envelope or tegument proteins of WSSV (Fig. 3J and 4H). Although PEN4 failed to interact with the test five viral structural proteins, PEN4 was able to interact with the outer surface of WSSV virion, which suggested that PEN4 could has the ability to bind to other structural proteins. To further confirm the above results, the phagocytic activity of hemocytes against WSSV was investigated. FITC was used to label the purified WSSV virion, and we observed that each recombinant penaeidin could significantly reduce the phagocytic activity of hemocytes against FITC-labeled WSSV virion (Fig. 6A). As shown in Fig. 6B, the phagocytosis rates of hemocytes in the treatment with WSSV by preincubation of rBigPEN-FL (22.53%), rBigPEN-PEN (26.07%), rPEN2 (23.97%), rPEN3 (24.03%) and rPEN4 (24.03%) were significantly reduced compared to that of the Trx tag control (33.20%) (P < 0.01) (Fig. 6B). In summary, these results convincingly suggested that penaeidins were able to inhibit WSSV entry into host cells by interacting with viral structural proteins.

**Figure 5.**
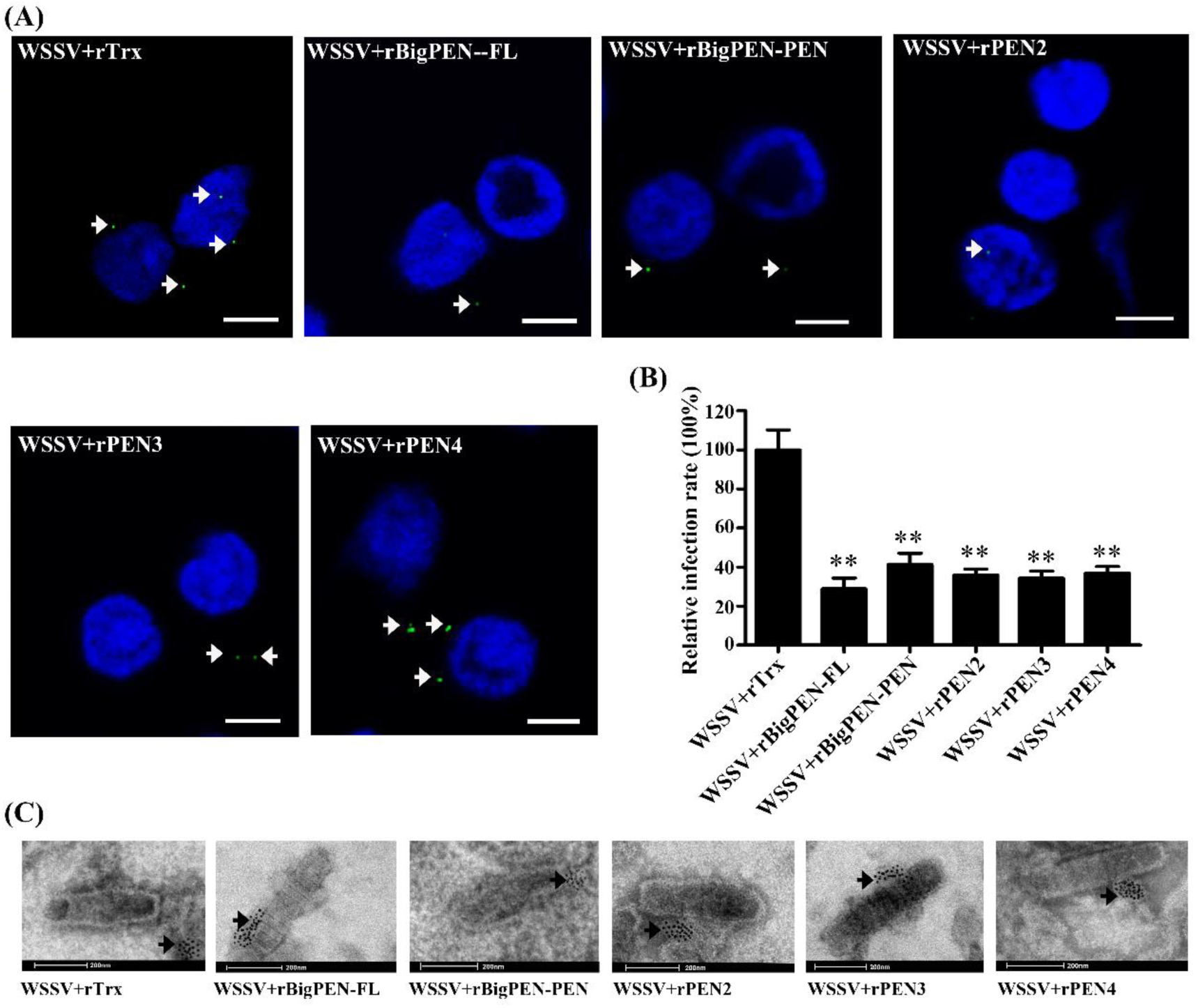
Penaeidins blocked WSSV infection *in vitro* and bound to the surface of WSSV virion. (A) Penaeidins blocked WSSV entry into hemocytes. Recombinant proteins BigPEN-FL, BigPEN-PEN, PEN2, PEN3 and PEN4 were firstly incubated with the FITC-labeled WSSV (green), and then added into the shrimp hemocytes. Subsequently, the hemocytes were stained with Hochest (blue) and observed under a fluorescence microscopy. Trx tag protein was used as a control. Scale bar, 25 μm. (B) Statistic analysis of WSSV infection-blocking rates of penaeidins corresponding to (A), and that of Trx tag protein was defined as 100%. (C) Recombinant penaeidins interacted with the surface of WSSV virion. The purified recombinant proteins BigPEN-FL, BigPEN-PEN, PEN2, PEN3 and PEN4 were firstly labeled with colloidal gold and then incubated with purified WSSV virion. After being stained with phosphotungstic acid, the viral suspension was adsorbed onto carboncoated nickel grids and observed under transmission electron microscopy (TEM). The Trx tag protein was used as a control; arrows showed locations of Trx, BigPEN-FL, BigPEN-PEN, PEN2, PEN3 or PEN4 labeled with colloidal gold. All experiments were performed three times with similar results.

**Figure 6.**
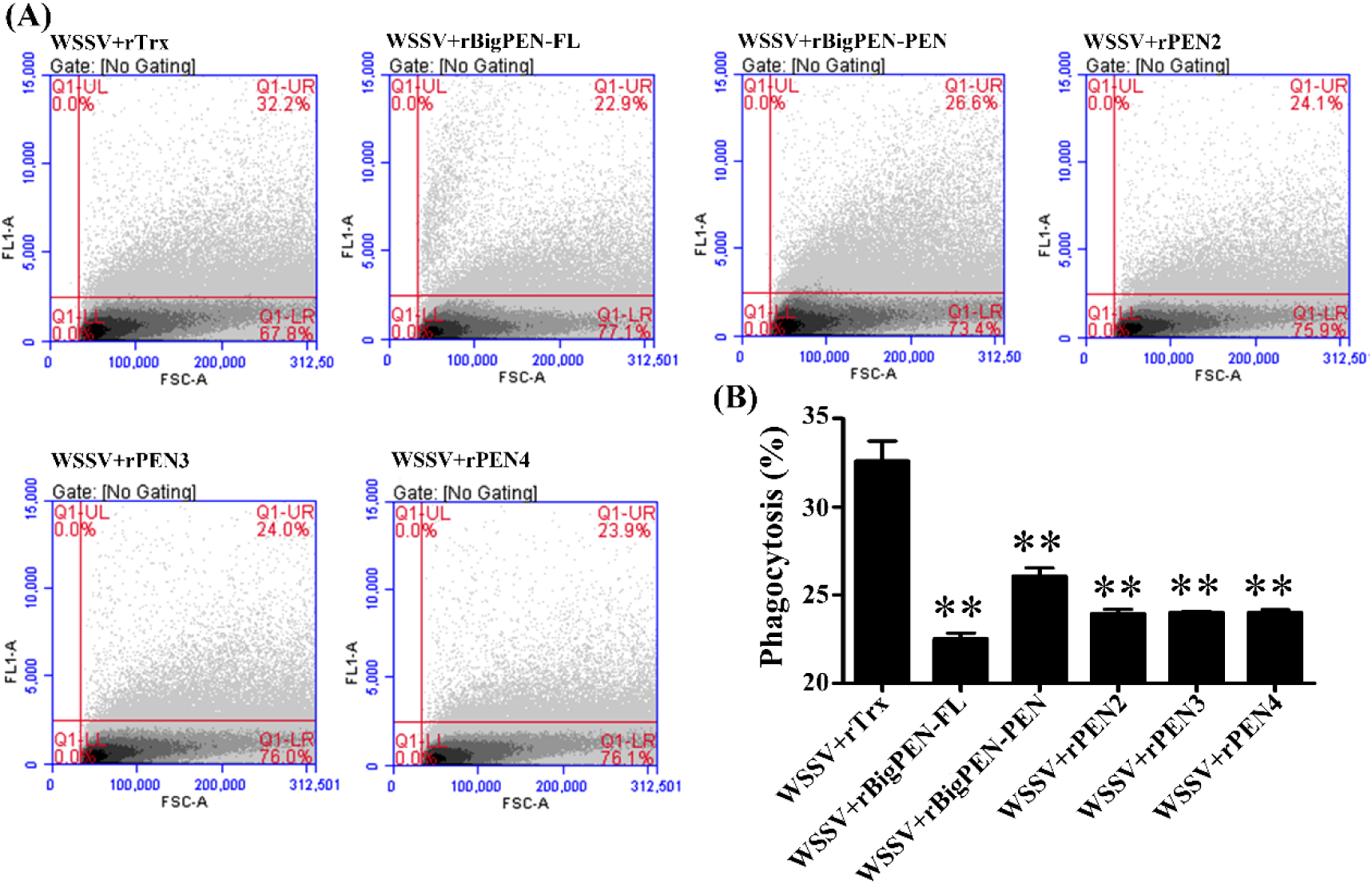
Effects of penaeidins on phagocytic activity of hemocytes against FITC-labeled WSSV virion. (A) Influences of recombinant proteins BigPEN-FL, BigPEN-PEN, PEN2, PEN3 and PEN4 or Trx (as a control) on phagocytic activity of hemocytes against FITC-labeled WSSV were detected by flow cytometry. Cells were examined by forward scatter (FSC, x-axis) and the phagocytosis of FITC-labeled WSSV was indicated by intracellular green fluorescence (y-axis) (^**^ *p* < 0.01). The scatter μlots represented one of the three flow cytometric detections. (B) Statistical analysis of phagocytic rates, Trx was used as a control. All the data were analyzed statistically by Student’s t-test (^**^ *p* < 0.01 and ^*^ *p* < 0.05). All experiments were performed three times with similar results.

### Penaeidins were regulated by conserved NF-κB pathway

In invertebrates, the transcriptional expression of AMPs was commonly regulated by conserved innate immune signaling pathways such as Toll, IMD and JAK-STAT pathways (29-32). In shrimp *L. vannamei*, Dorsal and Relish (NF-κB), the downstream transcription factors of Toll and IMD signaling pathways respectively, were regarded to be major factors to directly induce the production of AMPs in response to infection (33). To explore whether the expression of penaeidins were regulated by Dorsal and Relish, an RNAi *in vivo* experiment was performed. By qRT-PCR analysis, the mRNA levels of Dorsal and Relish could be effectively suppressed by corresponding dsRNAs (Fig. S1). We then observed that silencing of Dorsal or Relish resulted in varying degrees of downregulation in the transcript levels of BigPEN, PEN2, PEN3 and PEN4 under WSSV challenge *in vivo* (Fig. 7A). To address whether Dorsal and Relish were able to regulate the expression of penaeidins *in vitro*, Dual-Luciferase Reporter Assay and Electrophoretic Mobility Shift Assay (EMSA) were performed. We firstly obtained the promoter regions of four penaeidins including BigPEN, PEN2, PEN3 and PEN4 by Genome walking method, and then cloned them into the pGL3-Basic vectors, respectively. We observed that over-expression of *L. vannamei* Dorsal or Relish could significantly induce the promoter activities of all four penaeidins in *Drosophila* S2 cells (Fig. 7B). The above results suggested that both Dorsal and Relish were able to induce the expression of all four penaeidins *in vivo* and *in vitro*. Subsequently, BigPEN was chosen to be a representative to further confirmed the results in detail. We analyzed the 5^’^ flanking regulatory region of BigPEN, and found it contained two conserved κB motifs located at −349 to −339 (κB1, GTGTTTTTCGC) and −91 to −81 (κB2, GTGTTTTTTAC) respectively (Fig. 7C). Four vectors, including the wild type promoter region termed pGL3-κB12, and pGL3-κB-M1, pGL3-κB-M2 and pGL3-κB-M12 vector with a deletion mutant of one or both κB sites (Fig. 7C), were constructed to perform Dual-Luciferase Reporter Assay. We found that the promoter activities of pGL3-κB12, pGL3-κB-M1, pGL3-κB-M2 could be up-regulated by *L. vannamei* Dorsal over-expressed in S2 cells with 3.09-, 1.93-, 1.40-fold increases, whereas the pGL3-κB-M12 could not be any up-regulated (Fig. 7D). These results suggested that Dorsal could be able to interact with the conserved κB sites in the promoter region of BigPEN. To address this, EMSA was performed using purified 6His-tagged RHD domain of Dorsal protein (rDorsal-RHD) expressed in *E. coli* cells. As shown in Fig. 7E, *L. vannamei* rDorsal-RHD, but not the control rTrx, could effectively retard the mobility of the bio-labeled probe 1 (Line 5). We further observed that the DNA/Protein complex was faintly reduced by the competitive 2 × unlabeled probe 1, but markedly reduced by the competitive 100 × unlabeled probe 1 (Fig. 7E Line 6 and Line 7). In addition, rDorsal-RHD could not retard the mobility of the mutant bio-labeled probe 1 (Fig. 7E Line 3), which indicated the specificity of interaction between rDorsal-RHD and probe 1. Taken together, the results strongly suggested that NF-κB transcription factors (Dorsal and Relish) participated in the transcriptional expression of penaeidins in response to WSSV infection.

**Figure 7.**
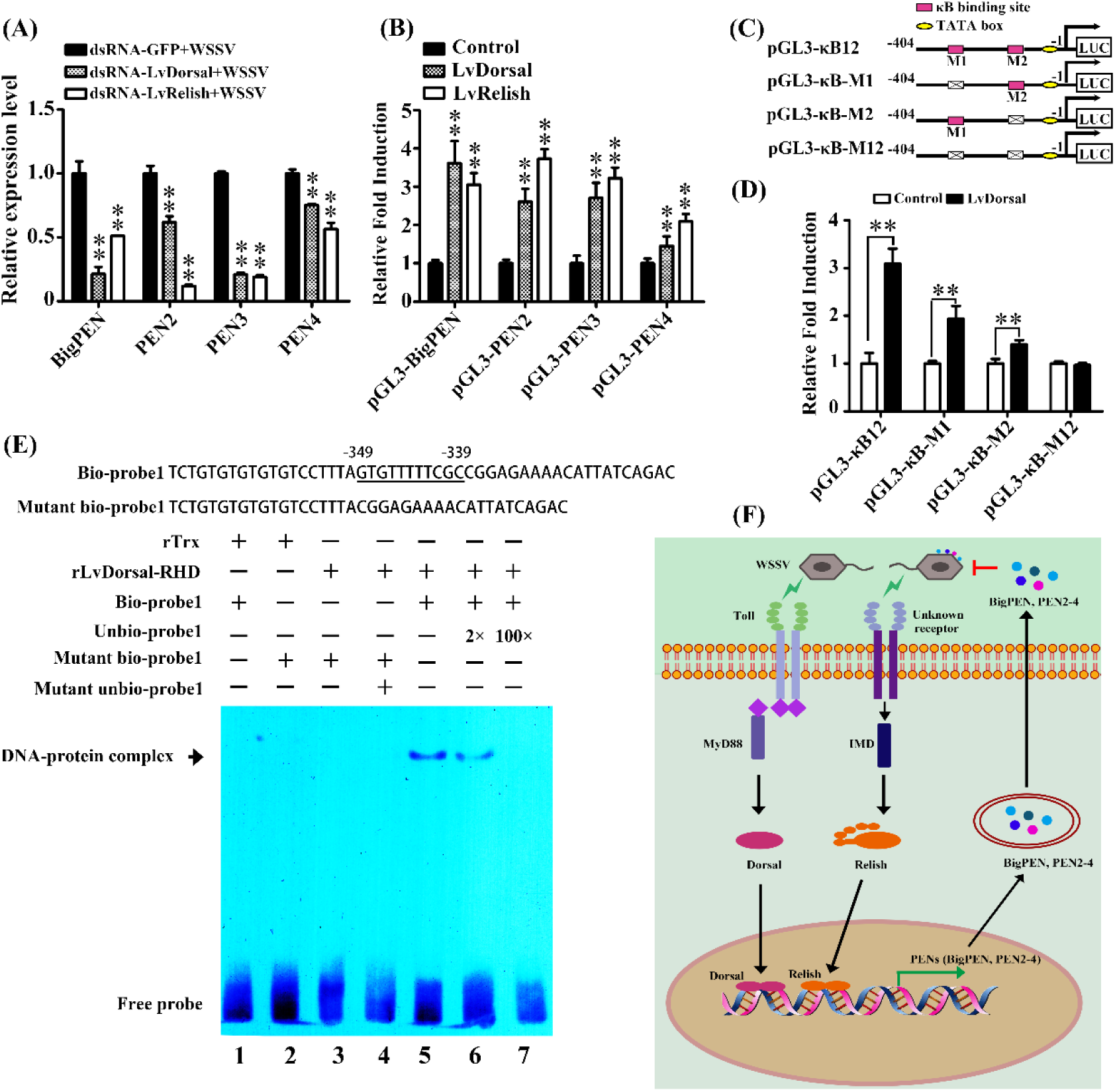
Penaeidins were regulated by NF-κB pathways. (A) The mRNA levels of BigPEN, PEN2, PEN3 and PEN4 in Dorsal- and Relish-silenced shrimps. The statistical significance was calculated using Student’s *t*-test, ^**^ *p* < 0.01 and ^*^ *p* < 0.05. (B) Dual-luciferase reporter assays were performed to analyze the effects of the over-expression of Dorsal and Relish on the promoter activities of BigPEN, PEN2, PEN3 and PEN4 in *Drosophila* S2 cells. All data were representative of three independent experiments. The value of the empty plasmid pAc5.1/V5-His A transfected cell was used as a control, which was set as 1.0. The bars indicated the mean ± SD of the relative luciferase activities (*n* = 3). The statistical significance was calculated using Student’s *t*-test (^**^ *p* < 0.01). (C) Schematic diagram of the promoter regions of BigPEN in the luciferase reporter gene constructs. (D) Dual-luciferase reporter assays were performed to analyze the effects of over-expression of Dorsal on the promoter activities of BigPEN with or deletion mutant of NF-κB binding motif (s). The bars indicated mean values ± S.D. of the luciferase activity (*n* = 3). Statistical significance was determined by Student’s *t*-test (^**^ *p* < 0.01). (E) Dorsal interacted with the NF-κB bind motif of BigPEN *in vitro*. EMSA was performed using biotin-labeled (Bio-) or unlabeled (Unbio-) probes containing or not containing the NF-κB binding motif of BigPEN. Biotin-labeled or mutated biotin-labeled double stranded DNA probes was incubated with 5 μg purified rDorsal-RHD protein, unlabeled probe was added to compete with binding, Trx protein was used as a control. All experiments were performed three times with similar results. (F) Model for penaeidins mediated antiviral mechanism against WSSV (See the text in detail).

## Discussion

In invertebrates, some antimicrobial peptides (AMPs) are identified as the viral responsive effectors, however the molecular mechanism underlying antiviral activities of AMPs is poorly understood. For example, two *Drosophila* AMPs, attC and dptB have been demonstrated to restrict Sindbis virus (SINV), but the actual action of their antiviral mechanism is still unknown (34). Herein, all four penaeidins found in *L. vannamei* including BigPEN, PEN2, PEN3 and PEN4, were chosen to explore their potential anti-WSSV activities. The important role of penaeidins during innate anti-WSSV response was established by using RNAi *in vivo* to silence each penaeidin expression. We observed that knockdown of each penaeidin resulted in higher viral loads, meanwhile each purified penaeidin could effectively confer shrimps against WSSV Moreover, we showed that the antiviral mechanism of penaeidins involved blocking viral internalization possibly because of their abilities to interact with outer surface of WSSV virion and its structural proteins. In summary, we, for the first time, provided some substantial evidences that penaeidins were a novel class of anti-WSSV effectors in shrimps.

The production of antiviral effectors represents the major host defense mechanism against viruses in invertebrates including shrimps, because they lack the adaptive immunity (35). Therefore, it is rationalized that identifying and characterizing novel antiviral molecules may shed important light into the innate immune antiviral response in shrimps. In this study, we focused our attention on penaeidins due to the following aspects: (i) penaeidins are a type of AMPs only found in penaeid shrimps; (ii) all four penaeidins from *L. vannamei* were significantly induced by WSSV infection (Fig. 1A); (iii) antiviral activities of penaeidins against WSSV have not been previously documented. In fact, similar to our observations, previous studies also showed that *L. vannamei* PEN2, PEN3 and PEN4 were strongly up-regulated at the early stage of WSSV infection (36), which could indicate that penaeidins played an innate antiviral response that acted early during infection to limit WSSV spread. For comprehensive analysis of the entire family of penaeidin during viral infection, we cloned another paralog and named it as BigPEN according to it containing additional RPT domain. All penaeidins from shrimps could be clustered into three subgroups, each subgroup contained one or two *L. vannamei* penaeidins. In specific, PEN3 located in subgroup 1, PEN2 and PEN4 located in subgroup 2, and BigPEN located in subgroup 3 (Fig. 1C). Considering the higher conservation of penaeidins in the same subgroup, the more similar function they might have. In that respect, the functions of *L. vannamei* BigPEN, PEN2, PEN3 and PEN4 during WSSV infection could be representative, to some extent, for those of penaeidins from other shrimps.

The name of penaeidins come from the originality of their structure and only found in penaeid shrimps (37). The penaeidins are composed of a conserved PEN domain including a proline-rich region (PRR) and a C-terminal region containing six cysteine residues (CRR) engaged in the formation of three intramolecular disulfide bridges (21). In this study, we identified a BigPEN, which contained an RPT domain prior to the PEN domain. It is apparent the RPT domain can’t interact with the test five viral proteins (Fig. 3H-3I) and the outer surface of WSSV virion (data not shown). However, the actual function of RPT domain is still unknown. In contrast, the PEN domain is conserved, and demonstrated to be able to interact with one or more viral structural proteins. Although PEN4 has failed to interact with the tested five viral proteins, it could be able to interact with other viral proteins as shown by that its capability to bind to outer surface of WSSV virion. This was further supported by that rPEN4, like other three penaeidins, can confer the hemocytes against WSSV entry.

The most probable scenario is that the antiviral activity of penaeidins is due to a direct interaction between the WSSV and penaeidins, and thereby inhibiting virus entry into target cells. Our current studies have provided several lines of evidence strongly support this notion. Firstly, all four penaeidins were able to interact with the outer surface of WSSV virion, and they, except PEN4, were shown to bind with one or more the tested five viral proteins. Secondly, the purified penaeidin proteins inhibited WSSV to enter hemocytes. Finally, the recombinant penaeidin proteins could significantly reduce the phagocytic activity of hemocytes against FITC-labeled WSSV virion. Indeed, interactions between structural proteins are common in the envelope viruses, and they might form complexes that have specific roles in host-viral interactions or the infectivity of viruses (14). Similar situation has also been observed in this envelope virus of WSSV. In this study, the identified penaeidins-binding viral proteins were the VP24, VP26, VP28 and VP16, among which VP24 and VP26 were the important tegument proteins, and VP28 and VP16 were the major envelope proteins of WSSV (38). VP28 was located on the outer surface of WSSV and involved in viral attachment to and penetration of shrimp cells (39). VP24 was a Chitin-binding protein and deemed to be a key factor involved in WSSV infection (40). VP26 was identified as an integral linker protein and can bind to host actin to help transport virions into host cells (41). In addition, VP24, VP26 and VP28 shared high sequence homology with each other, and they can form a complex termed ‘infectosome’ that has been regarded to be crucial to the infectivity of WSSV (42, 43). It is important to note that we are still unclear whether the interaction of penaeidins with viral proteins will interfere with the formation of ‘infectosome’ or other complexes involving host and viral proteins. Nevertheless, based on our results together with these previous observations, it is reasonable to conclude that the binding of penaeidins to viral tegument and envelope proteins attenuates WSSV infectivity, and inhibits WSSV internalization.

A common innate defense mechanism in invertebrates including shrimps is that immune signaling pathways, such as Toll, IMD and JAK/STAT pathways, regulate the production of some specific sets of effectors to defense against viral invasion (33). Identification of which pathway is responsible for the transcriptional expression of penaeidins in shrimps will help us better understand their mediated immune response to WSSV infection. Our results suggested that both of Dorsal and Relish (NF-κB), the downstream transcription factors of Toll and IMD pathways respectively, could be involved in the regulation of all four penaeidins after WSSV infection *in vivo*. This observation was further evidenced by that Dorsal was able to interact with the canonical κB motifs in the promoter region of BigPEN *in vitro* by EMSA assay. It is important to note that the one or both κB motifs could also be responsive to Relish, because that Relish strongly induced the expression of BigPEN was observed *in vitro* by Dual reporter genes assay (Fig. 7B). Similar situations are seen in other reports, for example, a κB motif in promoter of WSSV IE1 (wsv069) has been demonstrated as dual-responsive, that is, regulated by either Dorsal or Relish (44). In *Drosophila*, a single κB motif in the promoter of Metchnikowin (Mtk) has proved to bind both DIF and Relish (45). Additionally, other signaling pathways could be able to regulate the expression of some penaeidins, as shown by that the presence of several regulatory factors binding motifs, such as NF-κB, GATA, STAT and AP-1 in its promoter region (46). Thus, we proposed that the Toll and IMD signaling pathways, perhaps crosstalk with others, could work together in a collaborative manner to regulate the expression of penaeidins in response to WSSV infection. Such a regulatory pattern might be able to provide a rapid and tailored immune response against viral invasion according to varying degrees of severity.

In summary, we have identified penaeidins as a novel class of innate antiviral factors against WSSV for the first time. Based on our results, we proposed a model for the function of penaeidins in innate antiviral response (Fig. 7F). Infection of host cells with WSSV resulted in activation of Toll and IMD (NF-κB related) signaling pathway and production of penaeidins including BigPEN, PEN2, PEN3 and PEN4. These secreted penaeidins restricted WSSV infection by their interaction with the virion particles perhaps via viral structural proteins, which attenuated WSSV infectivity, and inhibited WSSV cellular entry. Future studies aimed at uncovering how penaeidins interact with viral proteins and how the resulting interactions effect viral internalization will lead to a better understanding of the innate immune antiviral function of penaeidins in shrimps.

## Materials and methods

### Animals and pathogens

Healthy *L. vannamei* (4 ~ 6 *g* weight each) were purchased from the local shrimp farm in Zhanjiang, Guangdong Province, China, and cultured in recirculating water tank system filled with air-pumped sea water with 2.5% salinity at 27 °C, and fed to satiation three times/ day on commercial diet. The Gram-negative *V. parahaemolyticus* were cultured in Luria broth (LB) medium overnight at 37 °C. Bacteria were quantified by counting the microbial colony-forming units (CFU) per milliliter on LB agar plates. The final injection concentration of *V. parahaemolyticus* should be controlled to yield ~1 × 10^5^ CFU/ 50 μl. WSSV was extracted from the WSSV-infected shrimp muscle tissue and stored at −80 °C. Before injection, muscle tissue was homogenized and prepared as WSSV inoculum with ~1 × 10^5^ copies in 50 μl PBS. In the pathogenic challenge experiments, each shrimp was received an intraperitoneal injection of 50 μl WSSV or *V. parahaemolyticus* solution at the second abdominal segment by a 1-ml syringe.

### RNA, genomic DNA extraction and cDNA synthesis

Total RNA was extracted from different tissues of shrimp using the RNeasy Mini kit (QIAGEN, Hilden, Germany). The genomic DNA shrimp tissues were extracted using a TIANGEN Marine Animals DNA Kit (TIANGEN, China), according to the manufacturer’s instructions. First strand cDNA synthesis was performed using a cDNA Synthesis Kit (Takara, Dalian, China), following the manufacturer’s instructions.

### cDNA cloning and sequence analysis

The partial cDNA sequence of BigPEN were obtained from transcriptomic sequencing of *L. vannamei* (26), and its full-length cDNA sequence was cloned by RACE-PCR according to previous method (47). The BigPEN, PEN2, PEN3 and PEN4 sequences were translated conceptually and the deduced protein was predicted using ExPASy (http://cn.expasy.org/). Similarity analysis was conducted using BLAST (http://blast.ncbi.nlm.nih.gov/Blast.cgi/) and the domain architecture prediction of the proteins was performed using SMART (http://smart.emblheidelberg.de). The neighbor-joining (NJ) phylogenic tree was constructed based on the deduced amino acid sequences of penaeidins protein by utilizing MEGA 5.0 software (48).

### Quantitative reverse transcription PCR

Quantitative reverse transcription PCR (qRT-PCR) was conducted to detect the mRNA levels of genes (penaeidins, Dorsal or Relish) for tissue distribution assay, pathogenic challenge experiments or silencing efficiency assay by RNAi *in vivo*. For tissue distribution assay, the shrimp tissues including eyestalk, epithelium, pyloric ceca, stomach, gill, heart, hepatopancreases, antenna, intestine, hemocytes were sampled. Three samples from each tissue were collected from 15 shrimps (5 shrimps pooled together as a sample). For pathogenic challenge experiments, 200 shrimps were divided into two experimental groups (100 shrimps in each group), in which each shrimp was injected with ~1 × 10^5^ CFU of *V. parahaemolyticus* or ~1 × 10^5^ copies of WSSV particles in 50 μl PBS, respectively. In addition, a group received with PBS injection was set as control. Hemocytes of challenged shrimps were collected at 0, 4, 8, 12, 24, 36 and 48 h post injection, and 3 samples at each time point were pooled from 9 shrimps (3 shrimps each sample). For silencing efficiency assay, shrimps were injected with corresponding dsRNAs (gene-specific dsRNA or the control GFP dsRNA). At 48 h post injection, three samples of hemocytes were collected from 15 shrimps (5 shrimps each sample). The method of total RNA extraction, cDNA synthesis and qRT-PCR analysis was performed as described (18). All samples were tested in triplicate. Primer sequences were listed in Table 1.

**Table 1.**
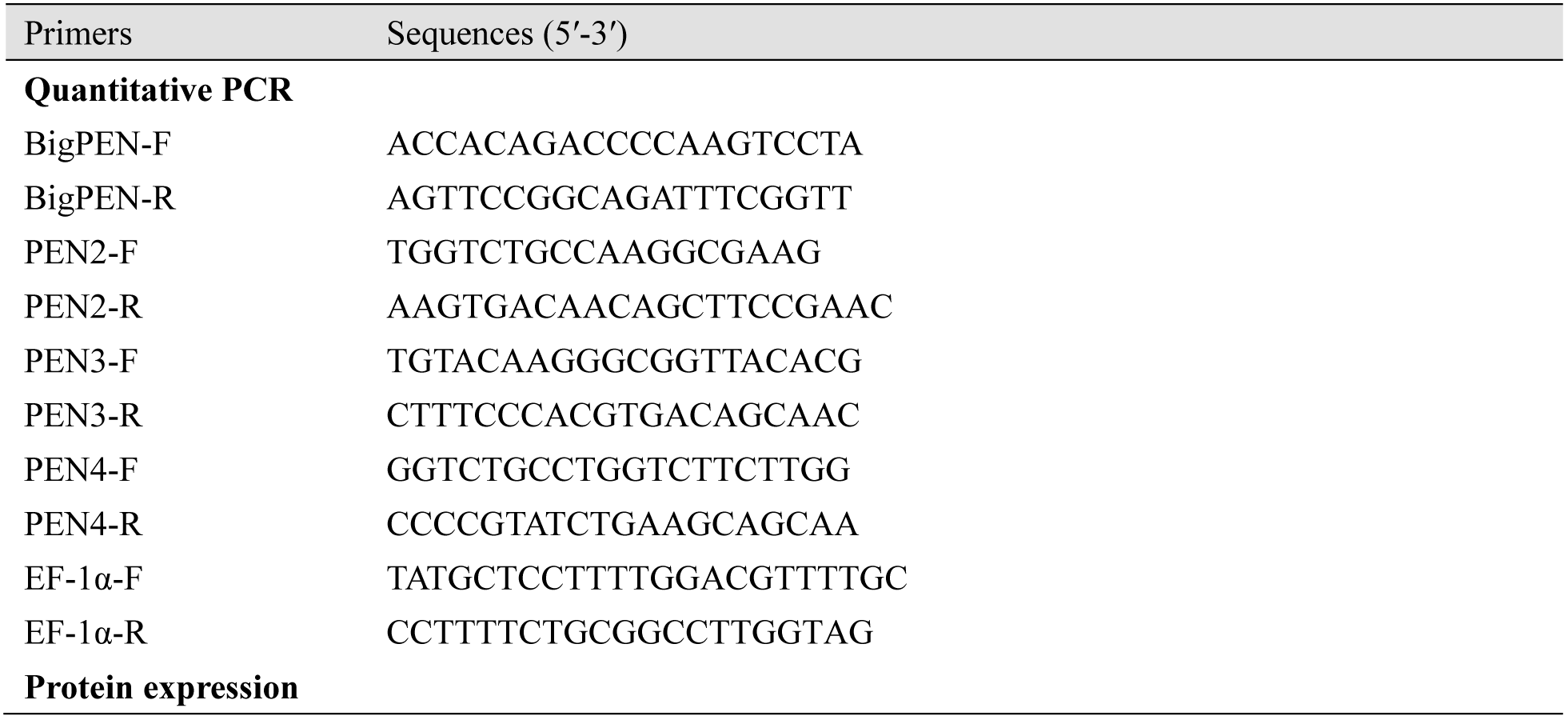
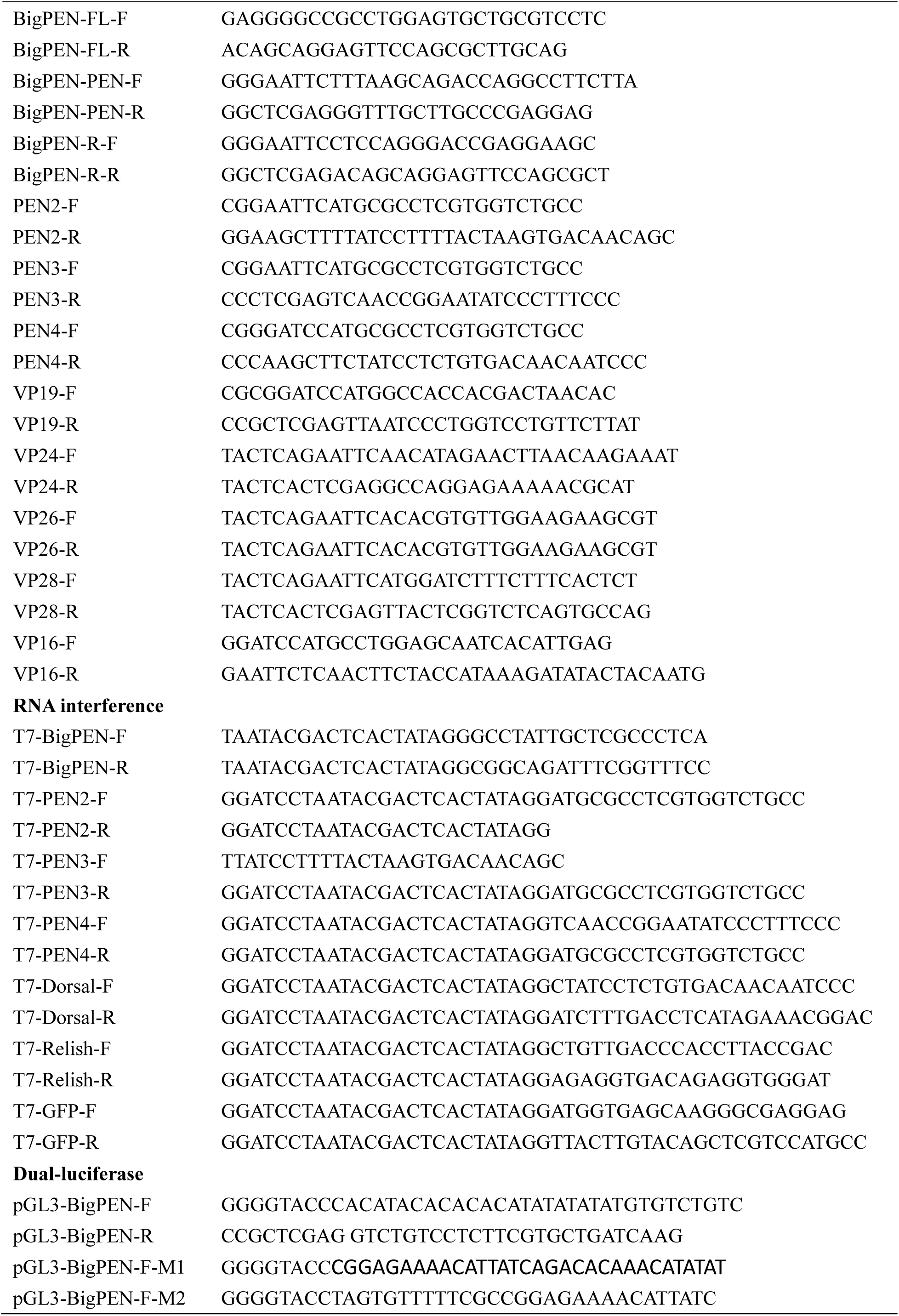
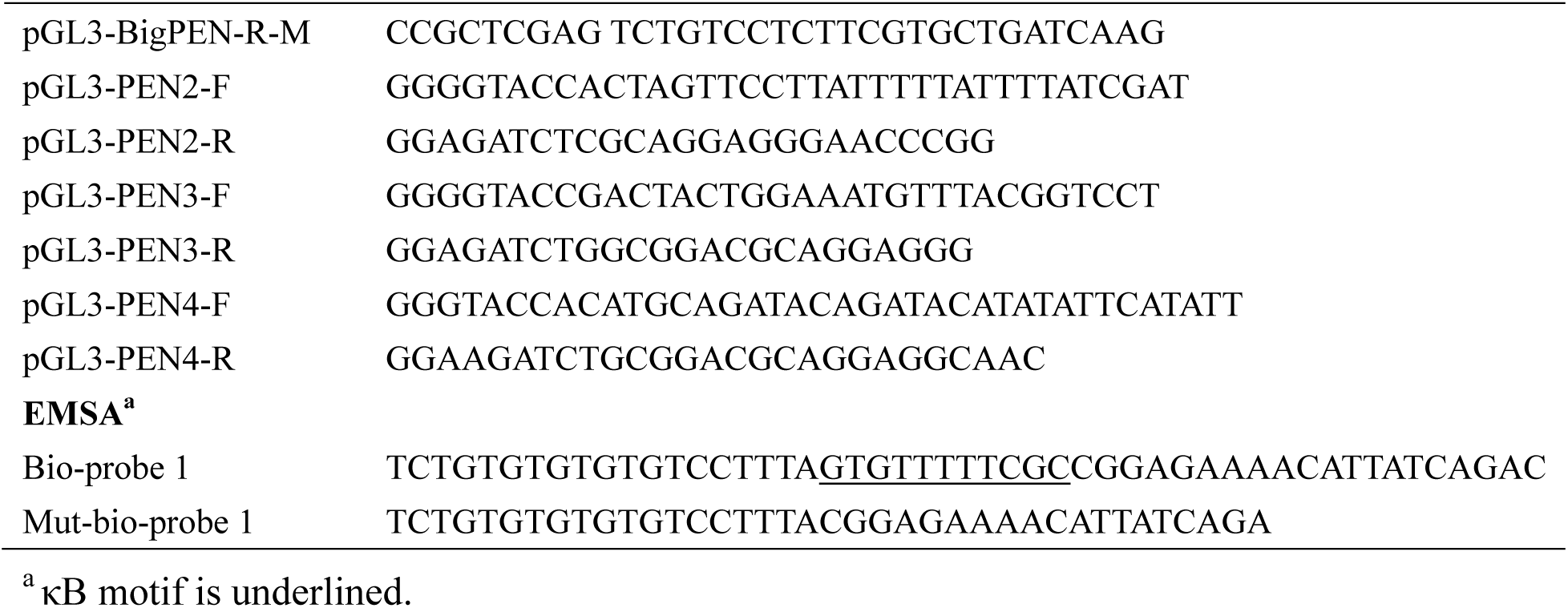
Primers used in this study.

### Recombinant proteins expression and purification

The open reading frames (ORFs) of penaeidins without the signal peptide coding sequences, were amplified by PCR using corresponding primers (Table 1) and subcloned into the pET-32a (+) plasmid (Merck Millipore, Germany) or pMAL-c2x plasmid (New England Biolabs, Ipswich, USA). Recombinant plasmids were transformed into Rosseta (DE3) cells for expression, and then the expressed proteins were purified by using Ni-NTA agarose (Qiagen, Germany) or Amylose Resin (New England Lab) according to the user’s manual. The purified proteins were checked by coomassie staining or western blotting. Concentration of the purified protein was determined using a BCA protein assay kit (Beyotime Biotechnology, China).

### Pull-down assay

Pull-down assays were performed to explore whether the recombinant full-length BigPEN (rBigPEN-FL), rBigPEN-PEN (C-terminal PEN domain of BigPEN), rBigPEN-R (N-terminal RPT domain of BigPEN), rPEN2, rPEN3 or rPEN4 could interact with the main structural proteins of WSSV (VP19, VP24, VP26, VP28 and VP16). These structural genes of WSSV were cloned into pGEX-4T-1 plasmid (GE Healthcare, USA) with specific primers (Table 1), expressed in Rosseta (DE3) *E. coli* strain, and purified with Pierce GST agarose (Thermo Scientific) recommended by user’s operation. For GST pull-down assays, 100 μl rBigPEN-FL (1 μg/ μl), rBigPEN-PEN (1 μg/ μl), rBigPEN-R (1 μg/ μl), rPEN2 (1 μg/ μl), rPEN3 (1 μg/ μl) or rPEN4 (1 μg/ μl) incubated with 100 μl GST tagged WSSV protein solutions (1 μg/ μl) at 4 °C for 1 h, respectively, and then the GST-bind resin was added and incubated for 2 h at 4 °C. The resin was washed with PBS thoroughly, the proteins were eluted with elution buffer (10 mM reduced glutathione and 50 mM Tris-HCl, pH 8.0) and then analyzed using 12.5% SDS-PAGE and western-blotting, the empty GST tag was used as control. For His pull-down assays, 100 μl rBigPEN-FL (1 μg/ μl), rBigPEN-C (1 μg/ μl), rBigPEN-R (1 μg/ μl) or rPEN3 (1 μg/ μl) incubated with 100 μl WSSV protein solutions (1 μg/ μl) at 4 °C for 1 h, respectively, and then the Ni-NTA bind resin was added and incubated for 2 h at 4 °C. The resin was washed with PBS thoroughly, the proteins were eluted with elution buffer (0.5 M NaCl, 20 mM Tris-Cl pH 7.4, and 300 mM imidazole), and then analyzed using 12.5% SDS-PAGE and western-blotting. For MBP pull-down assays, 100 μl rPEN2 and rPEN4 (1 μg/ μl) incubated with 100 μl WSSV protein solutions (1 μg/ μl) at 4 °C for 1 h, respectively, and then the MBP-bind resin was added and incubated for 2 h at 4 °C. The resin was washed with PBS thoroughly, the proteins were eluted with elution buffer (20 mM Tris-HCl pH 7.4, 0.2 M NaCl, 1 mM EDTA, 10 mM maltose), and then analyzed using 12.5% SDS-PAGE and western-blotting.

### Antiviral activities of recombinant proteins by RNA interference (RNAi) *in vivo* assay

The dsRNAs including BigPEN, PEN2, PEN3, PEN4 and GFP as a control were generated by *in vitro* transcription with T7 RiboMAX Express RNAi System kit (Promega, USA) using the primers shown in Table 1. The experimental groups were treated with the injections of dsRNA-BigPEN, dsRNA-PEN2, dsRNA-PEN3 or dsRNA-PEN4 (10 μg dsRNA each shrimp in 50 μl PBS), while the control groups were injected with equivalent dsRNA-GFP. Forty-eight hours later, the hemocytes from each group were sampled for qRT-PCR to detect the knockdown efficiency of BigPEN, PEN2, PEN3 and PEN4. Primer sequences were listed in Table 1.

To understand the function of penaeidins during WSSV infection, RNAi mediated knockdown of each penaeidin *in vivo* followed by viral titers and survival rates were checked. In WSSV challenge experiments, after 48 h post dsRNA injection, shrimps were injected again with 1 × 10^5^ copies of WSSV particles, and mock-challenged with PBS as a control. Then, shrimps were cultured in tanks with air-pumped circulatin seawater and fed with artificial diet three times a day at 5% of body weight for about 5 - 7 days following infection. After 48 h post WSSV infection, eight samples of muscle were collected. Muscle DNA was extracted with TIANGEN Marine Animals DNA Kit (TIANGEN, China) according to the user’s manual. The quantities of WSSV genome copies were measured by utilizing absolute quantitative PCR with primers WSSV32678-F/WSSV32753-R and a TaqMan fluorogenic probe as described previously (49). The survival rate of each group was recorded every 4 h. The Mantel-Cox (log-rank χ^2^ test) method was subjected to analyze differences between groups with the GraphPad Prism software.

In parallel, a series of rescue experiments were performed to monitor the effect of recombinant penaeidins on WSSV replication levels *in vivo* or survival rates in the knockdown of BigPEN, PEN2, PEN3 or PEN4 shrimps, respectively. Each recombinant protein of penaeidin (10 μg) were firstly incubated with WSSV for 1 h and then the mixture was inoculated into the experimental shrimps. The rMBP or rTrx proteins were used as controls. Likewise, viral loads and survival rates were analyzed as above.

### Infection-blocking assay *in vitro*

The virions were isolated from WSSV infected shrimps following the previous method (50). Intact WSSV particle was labeled with fluorescein isothiocyanate (FITC) (1 mg/ ml) for 2 h and then washed with PBS for three times. The FITC labeled WSSV were then mixed with rBigPEN-FL, rBigPEN-C, rPEN2, rPEN3, rPEN4 or rTrx (2 mg/ ml), respectively, and incubated at room temperature (RT) for 1 h. Hemocytes were collected from healthy *L. vannamei* by centrifugation (1000 *g*, 5 min) at RT and deposited onto a glass slide in 6-hole microtiter plates, and then 2 ml of the above virion suspension was added. Subsequently, the glass slices in the wells were washed with PBS three times, and fixed with 4% paraformaldehyde at RT for 5 min. After washed with PBS three times again, cells were incubated with Hochest 33258 (0.5 mg/ ml) for 10 min. Finally, the slices were visualized with confocal laser scanning microscope (Leica TCS-SP5, Germany). We calculated the colocalization quantities of FITC labeled WSSV with hemocytes manually from ten randomly selected visual regions, each includes at least 30 cells.

### Colloidal gold labeling and transmission electron microscopy

To investigate whether penaeidins could bind to WSSV virion, penaeidins were labeled with 10-nm-diameter gold nanoparticles (Sigma) following a previously reported method (51). Briefly, the pH of the colloidal gold was adjusted to at least 0.5 higher than the pI of each penaeidin by using 0.1 N HCL or 0.2 M Potassium carbonate. Then, the saturation isotherm was used to determine the protein/ gold ratio for the protein and colloidal gold. The minimal amount of protein necessary to stabilize the gold was determined by adding 1 ml of the colloidal gold to 0.1 ml of serial aqueous dilutions of the protein. Approximately 0.1 ml of colloidal gold was added to 100 μg of each penaeidin dissolved in 200 μl of PBS for 10 nM of gold. The solution was left to stand for 10 min, and then 1% polyethylene glycol (PEG) was added to a final concentration of 0.04%. The solution was left to stand for 30 min and centrifuged for 45 min at 50,000 × *g*. The supernatant was then removed, and the soft pellet was resuspended in 1.5 ml of PBS containing 0.04% PEG and stored at 4 °C. The colloidal gold-labeled penaeidin was diluted 1:10 in PBS containing 0.02% PEG. The empty Trx-His tag protein as a control was also labeled with gold nanoparticles. The purified virions were absorbed onto carbon-coated nickel grids, and incubated with labeled rBigPEN-FL, rBigPEN-PEN and rPENs (or rTrx) for 10 min at RT. After washing with distilled water three times, the samples were counterstained with 2% sodium phosphotungstate for 1 min and then observed under a transmission electron microscopy (JEM-100CXII).

### Phagocytic activity analysis

The phagocytic activity analyses were performed following the method modified from that of *Brousseau* et al (52). Briefly, hemocytes were collected from 50 healthy shrimps and washed with PBS triply and counted using a BD Accuri C6 Flow Cytometer (USA), and then mixed with 1 μg rBigPEN-FL, rBigPEN-PEN, rPEN2, rPEN3, rPEN4 or rTrx (as control) together with 100 μl FITC-labeled WSSV. After incubation at 28 °C for 1 h, hemocytes were detected using cytometry for the signals of FITC and the forward scatter (FSC) values of cells. A FSC threshold was determined through detection of free FITC-labeled WSSV to eliminate cell debris and WSSV, and the fluorescence boundary was set based on detection of the self fluorescences of untreated hemocytes. A total of 150,000 events were detected for each sample.

### Plasmid construction and Dual-luciferase reporter assay

The *L. vannamei* Dorsal and Relish expression vectors (pAc-LvDorsal-V5 and pAc-LvRelsih-V5) were obtained from our previous studies (53, 54). The reporter μlasmids including promoter regions of BigPEN, PEN2, PEN3 or PEN4 were cloned using primers (Table 1), and then linked into μgL3-Basic (Promega, USA) to generate μgL3-BigPEN (also named as μgL3-κB12), μgL3-PEN2, μgL3-PEN3 or μgL3-PEN4, respectively. Two putative NF-κB binding site (κB1, _-349_GTGTTTTTCGC_-339_ and κB2, _-91_GTGTTTTTTAC_-81_) in the promoter of BigPEN were predicted by JASPAR database (http://jaspardev.genereg.net/). The overlap extension PCR using primers (Table 1) was performed to construct three mutants of μgL3-κB12 with deletion of κB1 site, κB2 site or both sites, and named as μgL3-κB-M1, μgL3-κB-M2 and μgL3-κB-M12, respectively.

Given that no permanent shrimp cell line was available, *Drosophila Schneider* 2 (S2) cell line was used to detect the effects of *L. vannamei* NF-κB on promoters of *L. vannamei* BigPEN, PEN2, PEN3 and PEN4. S2 cells were cultured at 28 °C in Schneider’s Insect Medium (Sigma) containing 10% fetal bovine serum (Gibco). For dual-luciferase reporter assays, S2 cells were plated into a 96-well plate, at the next day, the cells of each well were transfected with 0.05 μg firefly luciferase reporter gene plasmids, 0.005 μg pRL-TK renilla luciferase plasmid (Promega, USA), and 0.05 μg proteins expression plasmids or empty pAc5.1A plasmids as controls. After 48 h transfection, the dual-luciferase reporter assays were performed to calculate the relative ratios of firefly and renilla luciferase activities using the Dual-Glo Luciferase Assay System kit (Promega, USA) according to the manufacturer’s instructions. All experiments were repeated for six times.

### EMSA assay

EMSA was performed using a Light Shift Chemiluminescent EMSA kit (Thermo) according to the method of previous study (55). Briefly, the biotin-labeled probe or unbiotin-labeled probe were designed the sequence containing the NF-κB binding motif sequence (GTGTTTTTCGC and GTGTTTTTTAC). The mutant probe was designed via deleting the NF-κB binding motif sequence. All the probes were synthesized by Life Technologies and sequences were listed in Table 1. EMSA was performed using a Light Shift Chemiluminescent EMSA kit (Thermo). The purified rDorsal-RHD (RHD domain of Dorsal) proteins (5 μg) were incubated with 20 fmol probes for the binding reactions between probes and proteins, separated by 5% native PAGE, transferred to positively charged nylon membranes (Roche), and cross-linked by UV light. Then the biotin-labeled DNA on the membrane were detected by chemiluminescence and developed on x-ray films, followed by enhanced chemiluminescence (ECL) visualization.

### Statistical analysis

All data were presented as means ± SD. Student *t*-test was used to calculate the comparisons between groups of numerical data. For survival rates, data were subjected to statistical analysis using GraphPad Prism software to generate the Kaplan–Meier μlot (log-rank χ^2^ test)

## Acknowledgement

This research was supported by National Natural Science Foundation of China (31772883); Tip-top Scientific and Technical Innovative Youth Talents of Guangdong special support program (No. 2016TQ03N504); Guangdong Natural Science Funds for Distinguished Young Scholars (2016A030306041); Fundamental Research Funds for the Central Universities (17lgpy62) and China Agriculture Research System (47). The funders had no role in study design, data collection and analysis, decision to publish, or preparation of the manuscript.

## Author Contributions

C.L., J.H. and B.X. designed all studies and analyzed all the data. C.L. wrote the manuscript. B.X., S.N. Q.F. and H.L. performed the experiments, with contributions by K.L., B.Y. S. W. and Q.F. to some of the vector constructions, quantitative PCR analyses, cell and shrimp culturing. All authors reviewed the manuscript.

---

**Figure S1.**
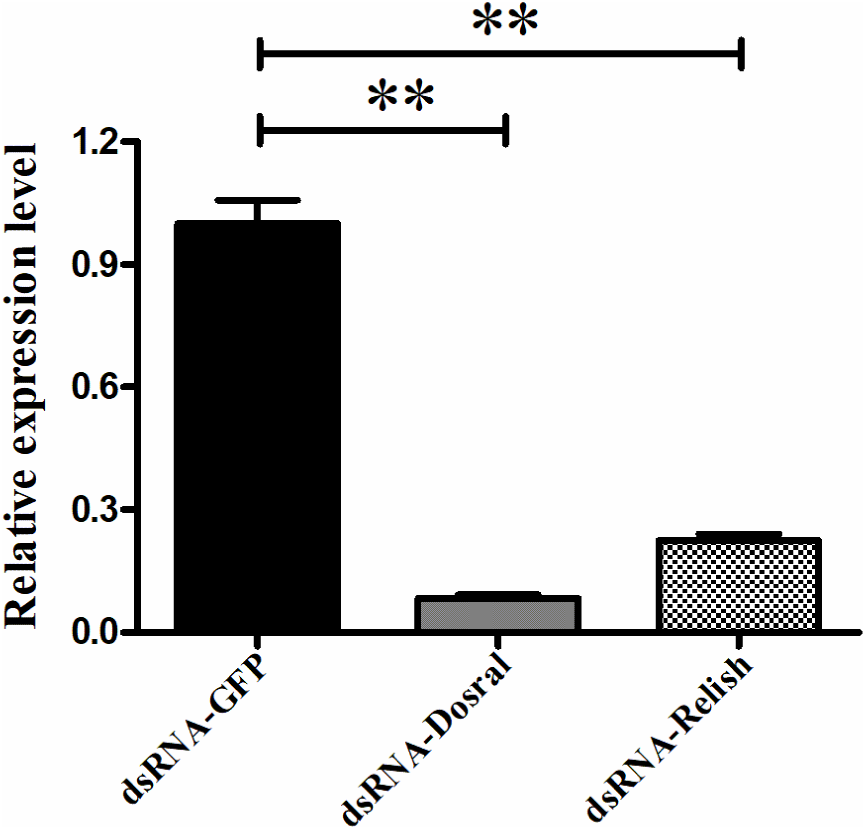
Effective knockdown for Dorsal and Relish in hemocytes by dsRNA was confirmed by qRT-PCR.

